# High-throughput protein target mapping enables accelerated bioactivity discovery for ToxCast and PFAS compounds

**DOI:** 10.1101/2025.03.20.644436

**Authors:** Diwen Yang, Xiaoyun Wang, Jiabao Liu, Yufeng Gong, Pranav Nair, Jianxian Sun, Xing Qian, Claire Cui, Hong Zeng, Aiping Dong, Rachel J. Harding, Nicola Burgess-Brown, Tyler S. Beyett, Datong Song, Henry Krause, Miriam L. Diamond, Derek L. Bolhuis, Nicholas G. Brown, Cheryl H. Arrowsmith, Aled M. Edwards, Levon Halabelian, Hui Peng

## Abstract

Chemical pollution is a global threat to human health, yet the toxicity mechanism of most contaminants remains unknown. Here, we applied an ultrahigh-throughput affinity-selection mass spectrometry (AS-MS) platform to systematically identify protein targets of prioritized chemical contaminants. After benchmarking the platform, we screened 50 human proteins against 481 prioritized chemicals, including 446 ToxCast chemicals and 35 per-and polyfluoroalkyl substances (PFAS). Among 24,050 interactions assessed, we discovered 35 novel interactions involving 14 proteins, with fatty acid-binding proteins (FABPs) emerging as the most ligandable protein family. Given this, we selected FABPs for further validation, which revealed a distinct PFAS binding pattern: legacy PFAS selectively bound to FABP1, whereas replacement compounds, PFECAs, unexpectedly interacted with all FABPs. X-ray crystallography further revealed that the ether group enhances molecular flexibility of alternative PFAS, to accommodate the binding pockets of FABPs. Our findings demonstrate that AS-MS is a robust platform for the discovery of novel protein targets beyond the scope of the ToxCast program and highlight the broader protein-binding spectrum of alternative PFAS as potential regrettable substitutes.

## Introduction

The global production and use of chemicals have expanded significantly, with over 350,000 chemicals currently registered worldwide^1^. Many of these chemicals ultimately enter human body, leading to undesired adverse health effects. In 2015, chemical contaminant exposure was estimated to contribute to approximately 9 million premature deaths globally^2^. However, the sheer scale of chemical contaminants far exceeds the capacity of traditional toxicity testing^3^, resulting in a limited understanding of their toxicity mechanism. This knowledge gap has often led to the widespread use of chemicals that were later proven to be hazardous and regrettable^4–6^. However, identification of the toxicity mechanism (*i.e.,* protein targets) of toxicants is extremely challenging and is the bottleneck for chemical management. Over the past decades, extensive efforts have been invested to assess the toxicity mechanism of chemical contaminants, by employing *in vitro* high-throughput screening (HTS) as a key strategy^7^.

The U. S. Environmental Protection Agency (EPA) launched the ToxCast program in 2007, which aims to systematically assess the molecular toxicity of ∼4,500 priority chemicals across ∼900 bioassays^8^. As the largest HTS campaign to date, the ToxCast program focuses on ∼300 protein targets, including common targets of toxins such as cytochromes and nuclear receptors (NRs). However, this subset of human proteins represents only 1.5% of ∼20,000 human proteins, leaving many potentially involved in pathways that contribute to toxicity unexplored. For instance, thalidomide was included in the ToxCast, yet its primary target cereblon (CRBN) identified in 2010^6^, was absent from the screening program. The broader expansion of the ToxCast screening protein targets has proven challenging due to the limited availability of bioassays for most human proteins. Indeed, traditional HTS assays are often designed specifically for one protein at a time, which do not scale across the full proteome. To address these gaps, it is important to develop new approaches that have the potential to screen protein targets at an unbiased, and potentially at a proteome-wide level.

Protein affinity selection mass spectrometry (AS-MS) is an unbiased approach for screening protein-ligand interactions^9^. Unlike traditional HTS bioassays, AS-MS is a direct binding assay that enables rapid and unbiased identification of protein-ligand interactions. Indeed, AS-MS has been successfully used to identify toxicants from environmental mixtures in our previous studies^10–14^, primarily focusing on a few proteins (*e.g.,* nuclear receptors). This study aimed to advance the AS-MS platform for scalable discovery of novel human protein targets for environmental chemicals, achieving an unprecedented throughput beyond traditional HTS bioassays. To test this, we decided to focus on 446 ToxCast chemicals and 35 perfluoroalkyl and polyfluoroalkyl substances (PFAS) that were prioritized by the U.S. EPA^15,16^. By employing a representative set of 50 human proteins not covered in the ToxCast program, we assessed 24,050 interactions within three weeks, leading to the identification of 35 novel protein-toxicant interactions. Among these, fatty acid-binding proteins (FABPs) emerged as the most ligandable protein family were selected for further validation. Notably, legacy PFAS selectively bound to FABP1, while alternative PFAS interacted with all seven FABP isoforms.

## Results

### Benchmarking the AS-MS platform

The AS-MS platform adapts our previous affinity purification with nontargeted analysis (APNA) method^10–14^, as illustrated in Fig. 1a. In brief, all prioritized chemicals including 446 ToxCast compounds and 35 PFAS (Table S5) were mixed in one pool for AS-MS screening to increase the throughput. 100 nM of pooled compounds were incubated with 1 μM His-tagged proteins and Ni-NTA magnetic beads in 96-well plate format. The ligands binding to target proteins were isolated from free chemicals and proteins through affinity purifications for liquid chromatography–mass spectrometry (LC–MS) analysis. Putative ligands for each protein were identified by comparing the LC-MS signals (*i.e.,* fold change > 10, *p* < 0.05) against other proteins in the same batch. The ligands were further confirmed by orthogonal assays, such as the fluorescence displacement assay or protein-ligand co-crystal structure by X-ray crystallography.

**Fig. 1.**
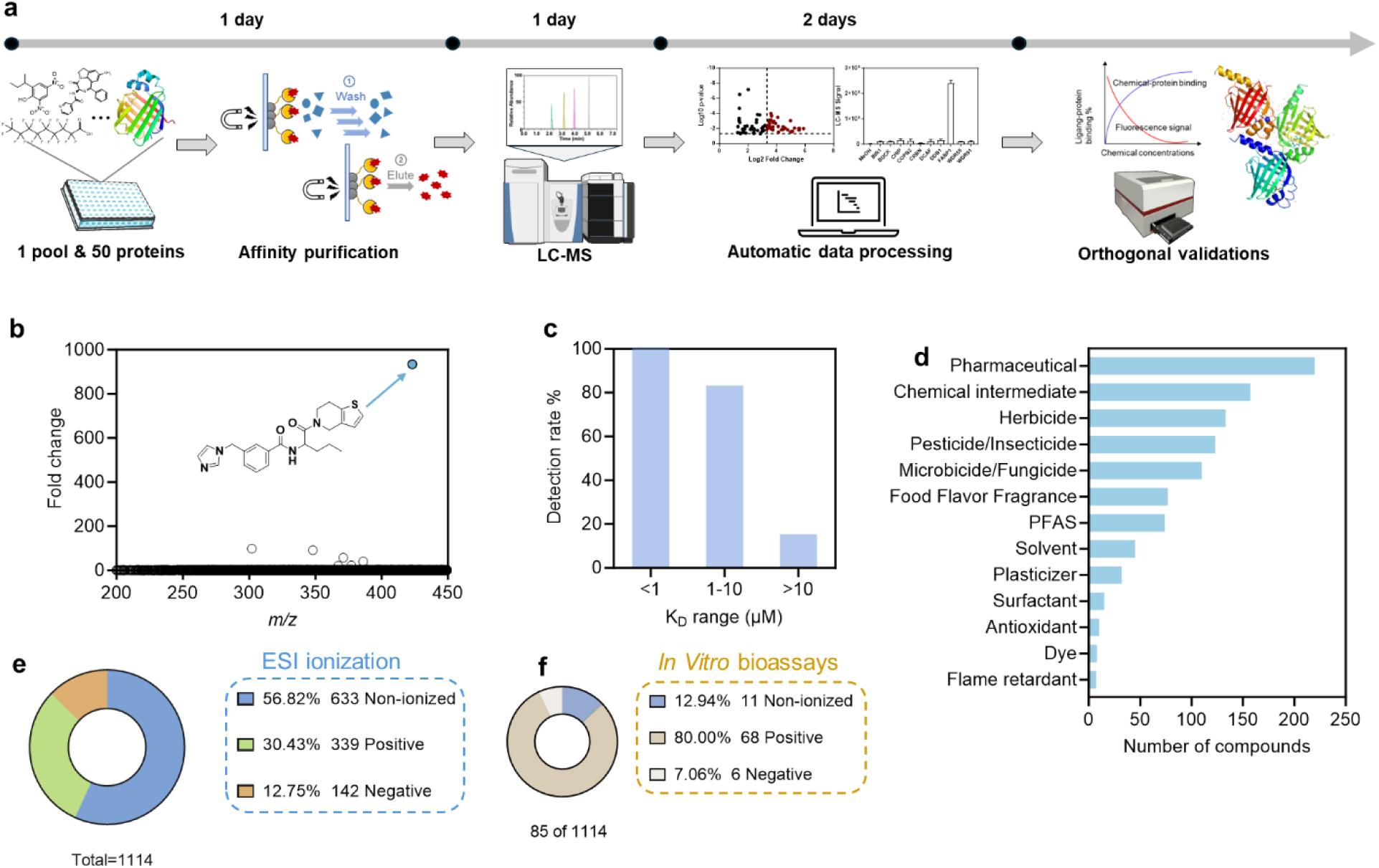
Benchmarking the AS-MS platform for scalable protein target discovery. (**a**) Workflow of AS-MS platform. (**b**) Benchmarking the AS-MS platform using a double-blind experiment. The AS-MS platform successfully identified a known ligand (blue dot, K_D_ = 60 nM) of WD repeat-containing protein 5 (WDR5) from a library of ∼16,000 compounds. (**c**) Detection rate of compounds according to their dissociation constants (K_D_). (**d**) Top 10 compound categories in the 1,114-chemical library based on the Chemical Products Categories (CPCat) database^68,69^. (**e**) The fraction of MS activity compounds in the 1,114-chemical library. (**f**) A subset of 85 bioactive compounds from the library exhibited AC₅₀ < 10 µM against nuclear receptors.

To pilot the AS-MS platform for large-scale screening, we first conducted a double-blind experiment by spiking a known ligand (K_D_ = 60 nM) of WDR5 protein to ∼16,000 compounds. This known compound was identified as the top hit within one week, with a high enrichment fold of 934 compared to negative control proteins (Fig. 1b). Encouraged by the promising results, we further benchmarked the platform using eight human proteins (WDR5, WDR91, HDAC6, LRRK2, USP21, CRBN, DCAF1, and GID4) representing diverse biological functions were selected. A total of 101 verified ligands for these proteins were mixed as a chemical pool, to evaluate the sensitivity and selectivity of our AS-MS platform. The high selectivity of the assay was demonstrated by the strong affinity ligands (K_D_ < 1 µM) detected for WDR5, GID4, DCAF, CRBN, and HDAC6, which exhibited high fold changes while showing little to no binding to other proteins (Fig. S1). Collectively, 100%, 83%, and 15% of strong-(K_D_ < 1 µM), moderate (1 µM < K_D_ < 10 µM) and weak-affinity (K_D_ > 10 µM) ligands were detected by our AS-MS method, respectively (Fig. 1c). Strong-affinity binding in the nanomolar (nM) range is typically essential for environmental contaminants to cause adverse health effects (*e.g.,* K_D_ = 1 nM for dioxins on AhR^17^), at environmentally relevant concentrations. Therefore, the high sensitivity (100%) of our AS-MS platform for strong-affinity ligands is well suited for protein target profiling of environmental toxicants.

After benchmarking the AS-MS platform, we proceeded to design a library of prioritized chemicals for screening. We decided to focus on the ToxCast (1,040 compounds) and PFAS (74 compounds) chemical libraries prioritized by the U.S. EPA, as they represent a broad spectrum of commercially used substances with widespread human exposure^15,16,18,19^. These compounds span more than ten categories, including pharmaceuticals and pesticides, among others (Fig. 1d). PFAS compounds were selected as they are well-known ‘forever chemicals’ with concerns regarding their high persistence, bioaccumulation and toxicity^20–22^. The U.S. EPA recently established a library of 74 PFAS for toxicity research according to a chemical category-based prioritization approach^16,23^. As high-resolution MS is used for detecting isolated ligands, we further prioritize compounds with high LC-MS sensitivity for AS-MS screening. Among 1,114 compounds tested, 481 compounds (43.2 %), including 446 ToxCast compounds and 35 PFAS, were detectable by LC-MS at 100 nM (Fig. 1e). To assess the potential coverage of AS-MS for bioactive compound screening, we compared the LC-MS detection of 1,114 compounds to their reported bioactivity towards nuclear receptors^24^. Among them, 85 compounds showed strong bioactivity with AC_50_ below 10 µM (Fig. 1f), while the majority were inactive. Notably, 74 of the 85 potent compounds (87.1%) were detectable by LC-MS, demonstrating that most bioactive toxicants are LC-MS active and are covered by AS-MS. This might not be surprising, because H-bonds are typically important for both protein binding and LC-MS ionization^25,26^. In total, 481 LC-MS active compounds were selected for AS-MS screening in subsequent studies.

### Application of the AS-MS platform to 481 ToxCast/PFAS chemicals and 50 human proteins

The benchmarked AS-MS platform was then applied to 50 human proteins (Table S3) to discover novel targets of the select 481 compounds. We selected proteins with a range of molecular weights, protein structures and biological functions associated with various diseases (Fig. 2a). While these proteins have been proposed as important targets for disease treatment and drug development,^27,28^ none of these proteins have been previously assessed in the ToxCast program, probably due to the lack of bioassays. For instance, as mentioned earlier, CRBN was identified as the primary target of thalidomide in 2010,^6^ three years after the launch of the ToxCast program. Human fatty acid-binding proteins (FABP1-7) are another protein family of interests, primarily responsible for the transport of hydrophobic ligands, such as fatty acids, within cells^29^. Among these, FABP1 has been thoroughly studied for its role in regulating the toxicokinetic of PFAS^17,22,23^,. Additionally, leucine-rich repeat kinase 2 (LRRK2), a protein associated with Parkinson’s disease (PD), has been the subject of over 3,000 publications, with three ligands progressed to preclinical trails^32,33^.

**Fig. 2.**
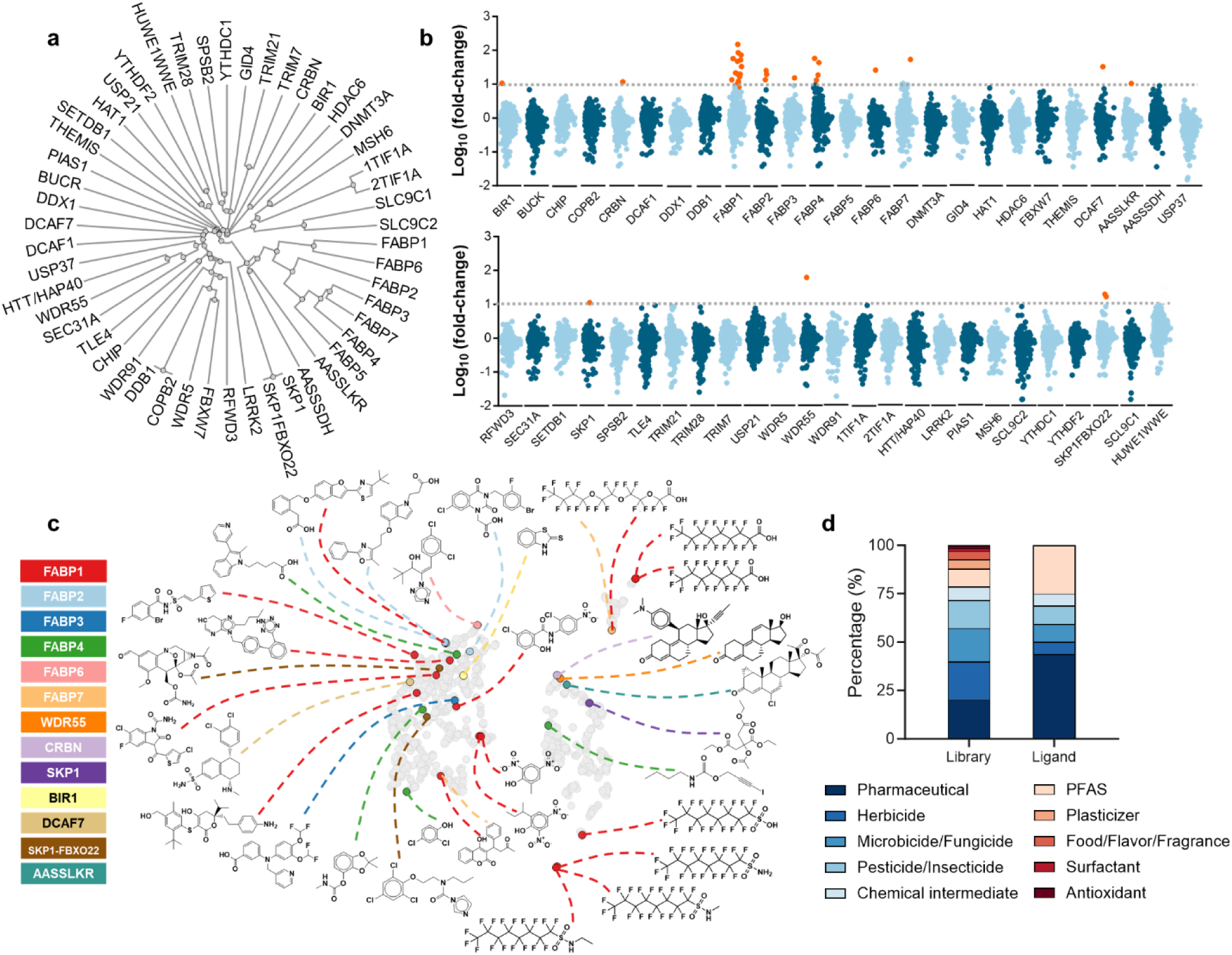
Mapping interactions between 481 selected chemicals and 50 human proteins. (**a**) Overview of 50 selected human proteins. (**b**) Scatter plot of hits interacting with 50 proteins. The gray line represents a fold change cutoff of 10. Red dots indicate hits with a fold change greater than 10 and a *p-value* less than 0.05. The fold change for each compound-protein interaction was calculated by dividing the average LC-MS peak intensity of the target protein by the average signal across all other proteins on the plate and *p-values* were determined using Student’s t-tests. (**c**) Chemical space of the 1,114 compounds clustered based on their molecular fingerprints and visualized using uniform manifold approximation and projection (UMAP). Detected ligands are highlighted in different colors according to their target proteins, with chemical structures labeled on the plot. (**d**) Comparison of the top 10 compound categories in the 481 selected compounds (left column) with the categories of ligands that showed protein interactions in the AS-MS screening (right column).

The screens were completed within three weeks, assessing 24,050 interactions (50 proteins × 481 compounds). We identified 35 compound-protein pairs targeting 14 proteins, representing a 10-fold enrichment compared to other proteins screened in the same batch (Fig. 2b). This leads to an average hit rate of 0.15% at the compound level which is 2.9-times lower than the hit rate of NRs covered by the ToxCast program (0.44%)^24^. This is expected, as many proteins including NRs covered by the ToxCast program, are among the most studied and ligandable proteins. Among the 50 proteins studied here, FABPs are the most detected targets, with 1-16 putative ligands discovered for each of FABP1-7 (Fig. 2c). This might not be surprising as FABPs are major transporters of fatty acids, and they have been reported to bind to xenobiotics including ‘forever compounds’ PFAS^25,30,31,34–39^. Among 35 compound-protein interactions, 8 interactions have been reported in previous studies, such as the binding of perfluorooctanesulfonic acid (PFOS) to FABP1^36^. The detected ligands showed diverse structures distributed across the chemical space of compound library (Fig. 2c). These include clusters of compounds with 21 chemical scaffolds such as multiple aromatic rings, fluorinated compounds, and those with primarily aliphatic structures (Fig. S3). Additionally, these ligands were primarily enriched in pharmaceuticals and PFAS (Fig. 2d), highlighting their potential health concerns targeting these novel protein targets.

### Discovery of ligands with new chemotypes binding to human FABPs

Among 14 novel protein targets, FABPs are of particular interest, not only due to the number of ligands but also their high AS-MS enrichment fold. As a result, we decided to focus on FABPs and used an orthogonal fluorescence displacement method^25^ to validate the bindings. Among the 26 interactions for FABPs detected by AS-MS, five compounds with background fluorescence were excluded. Among the remaining interactions, 16 were successfully validated by the fluorescence displacement method, with K_D_ ranging from 0.02 to 149 μM, leading to a validation rate of 76.2%. The lack of validation for 5 interactions was probably due to two reasons. 1) Hydrophobic compounds with low water solubility might not be validated by the fluorescence assay; 2) Some chemicals might be degraded during the storage in DMSO at-80 °C, leading to overestimated concentrations for validation experiments. Thus, the validation rate of AS-MS results might be underestimated. However, the limitations of fluorescence displacement assays, which are common in traditional HTS bioassays, highlight key advantages of AS-MS: 1) the high sensitivity of MS enables the use of very low chemical concentrations, eliminating solubility issues; and 2) AS-MS is not sensitive to absolute chemical concentrations, and thus potential degradation in large chemical libraries during storage does not impact screening. These additional strengths make AS-MS well-suited for protein target discovery.

Among seven FABPs, more ligands were detected for FABP1 (15) and FABP6 (5), albeit with lower binding affinities (K_D_ between 0.05-9.6 µM and 3.32-149 µM, respectively) compared to the other FABPs (Fig. 3a). Principal component analysis (PCA) further revealed that FABP1 and FABP6 cluster closely based on their ligand interaction patterns, suggesting similarities in their binding profiles (Fig. 3b). Given these similarities, we investigated whether the accessibility of compounds to protein binding pockets plays a key role in determining ligand interactions^40^. To test this, the solvent-accessible volumes of seven FABP isoforms in their ligand binding conformations were calculated using the CASTp (Fig. 3c)^41^. Among the seven FABPs, the binding cavity sizes of FABP1 and FABP6 were calculated as 361 Å^3^ and 393 Å^3^, respectively, bigger than the five other FABPs (268-304 Å^3^). This may explain the larger number of ligands detected for FABP1 and FABP6, as their larger binding pockets can accommodate more ligands. Further PCA analysis showed that PFOS and perfluorononanoic acid (PFNA) clustered closely with FABP1, indicating a selective binding of these PFAS to FABP1 (Fig. 3b). Due to their structural similarity to fatty acids, it is expected that PFAS would be prominent among the detected ligands, binding to FABP1 in the low micromolar range^25,30,31,34–39^. This is consistent with FABP1’s primary role as a transporter of endogenous fatty acids in the liver. For example, PFOS and perfluorooctanoic acid (PFOA) were found to bind to FABP1 with K_D_ of 8.10 and 18.8 μM, respectively^25^. However, the other novel ligands of FABP1 discovered by AS-MS showed very different chemotypes compared to known ligands with alkyl chains. Indeed, eight of 16 FABP1 ligands are aromatic compounds showing even stronger binding affinities than alkyl ligands, such as FK-739 (K_D_ = 0.56 µM) and DTXCID5027304 (K_D_ = 4.45 µM) (Fig. 3d).

**Fig. 3.**
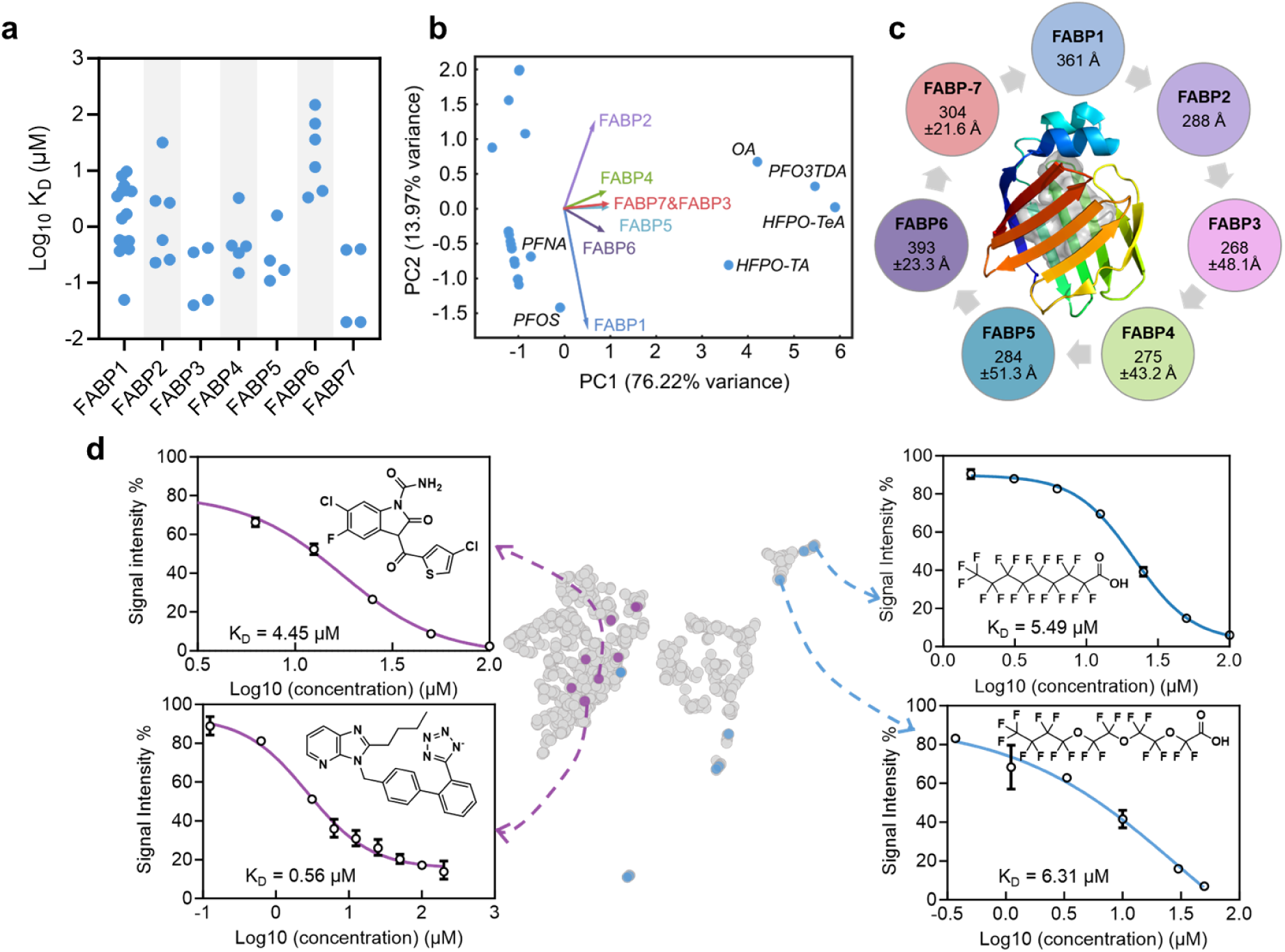
Distinct and novel interactions between ToxCast/PFAS chemicals and FABPs. (**a**) Overview of the binding affinity range (Log_10_ K_D_, µM) for ligands interacting with seven FABPs measured by fluorescence displacement assay. Each blue dot represents a single ligand. (**b**) Principal component analysis (PCA) of ligand binding affinities (Log10 K_D_, µM) across seven FABPs. Fluorescence displacement assays were used to determine binding affinities, with 1,8-ANS as the probe for FABP1, 2, 3, 4, 5, and 7, and bisANS for FABP6, while oleic acid (OA) K_D_ values for the seven FABPs were obtained from the literature^29,66,67^. (**c**) Solvent-accessible volumes (light gray) of FABPs. FABP1 (PDB: 3STM) is shown as a representative structure for the seven FABPs. For FABP isoforms with multiple available crystal structures, mean and standard deviation values were calculated to determine solvent-accessible volume variations. (**d**) Distribution of known ligands (blue dots) and newly discovered ligands (purple dots) for FABP1 within the chemical space of the 1,040 ToxCast compounds and 74 PFAS compounds. Dose-response results from the fluorescence displacement assay validation are shown, along with the measured dissociation constants (K_D_). DTXCID5027304 (CAS, 135080-03-4; top left panel) was tested with six concentrations (3.12, 6.25, 12.5, 25, 50 100 µM); FK-739 (CAS, 136042-19-8; bottom left panel) was tested with nine concentrations (0.12, 0.62, 3.12, 6.24, 12.5, 25, 50, 100, 200 µM); PFNA (CAS, 375-95-1; top right panel) was tested with seven concentrations (1.78, 3.12, 6.25, 12.5, 25, 50 100 µM); PFO3TDA (CAS, 330562-41-9; bottom right panel) was tested with six concentrations (0.37, 1.11, 3.33, 10, 30, 50 µM).

### Discovery of structure-specific binding of legacy and alternative PFAS to FABPs

Building on our findings of diverse compound classes targeting FABPs, we observed distinct binding patterns among different PFAS groups. Both legacy PFAS and perfluoroether carboxylic acids (PFECAs) were enriched by FABPs in AS-MS, yet their interactions varied. Due to the environmental persistence, bioaccumulation, and toxicity^42–46^, legacy PFAS (commonly known as “forever chemicals”) have been phased out and replaced with alternative PFAS^47–50^. Among these alternatives, PFECAs were designed as replacement for legacy PFAS with the similar structures^51^ (see representative structures in Fig. 4a). Notably, while legacy PFAS were only enriched by FABP1, PFECAs (*e.g.,* PFO3TDA) were enriched by other FABPs in addition to FABP1. Inspired by this observation, we systematically assessed the FABP binding of six legacy and six alternative PFAS (Fig. 4a). Consistent with the AS-MS results, the fluorescence displacement assay revealed that legacy PFAS strongly bind to FABP1, while showing minimal binding to other FABPs (Fig. 4b). In contrast, long-chain PFECAs bound to all seven FABPs with greater affinities than legacy PFAS. For instance, PFO3TDA, previously identified as an FABP1 ligand^25^, strongly bound to all seven FABPs with K_D_ values ranging between 0.02 and 4.35 µM (Table S1). Similarly, HFPO-TeA and HFPO-TA bound to all seven FABP isoforms with the K_D_ values ranging from 0.02-3.32 µM and 0.25-69.0 µM, respectively (Fig. 4c and Fig. S5). In contrast, shorter-chain PFECAs showed no detectable FABP binding (Fig. S5), underscoring the importance of the hydrophobic interactions of fluorocarbon tails^25,31^. Collectively, despite their structural similarities, legacy and alternative PFAS showed structure-specific binding patterns to FABPs.

**Fig. 4.**
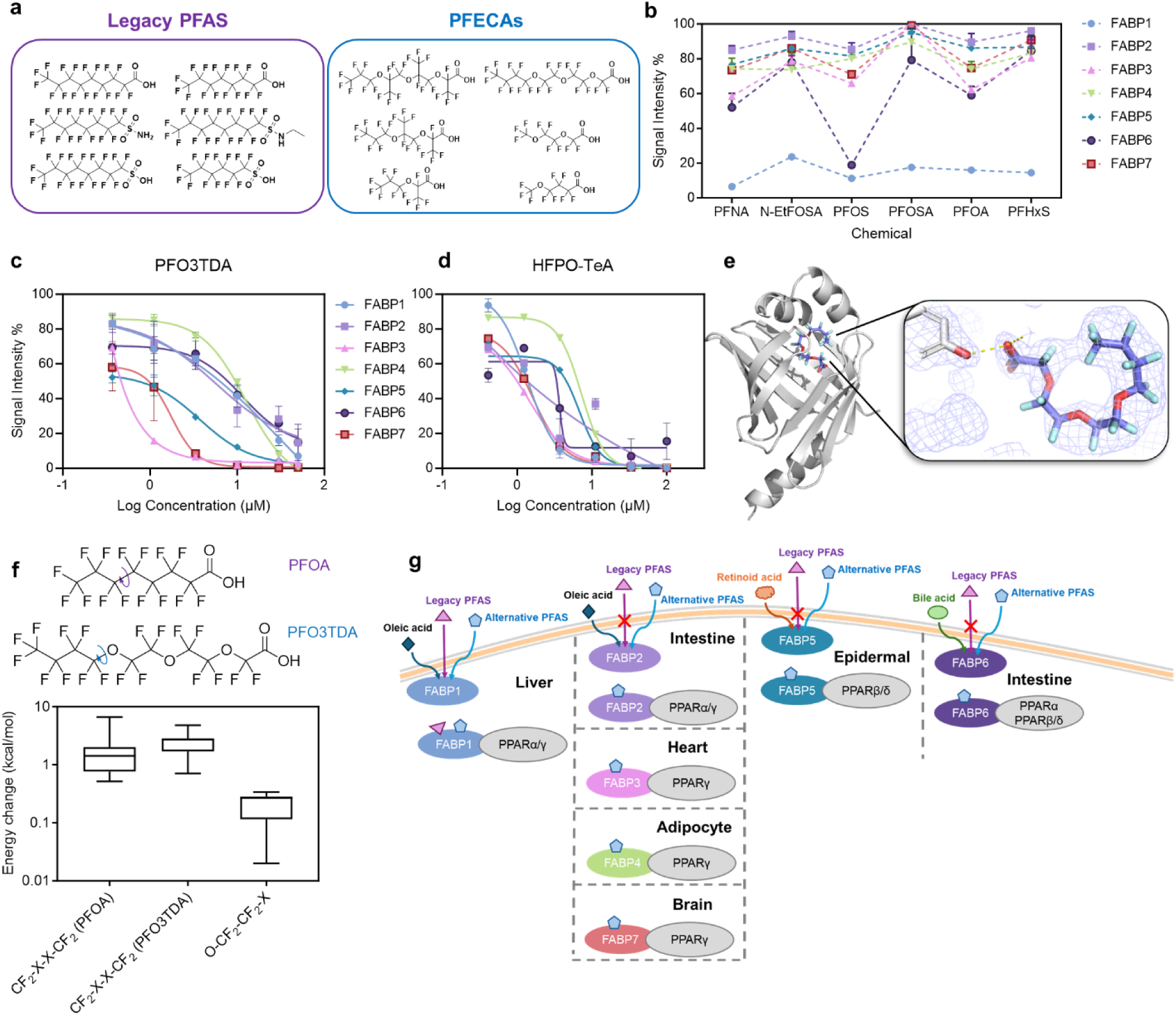
Selective interactions between alternative PFAS and FABPs. (**a**) Chemical structures of representative legacy PFAS and PFECAs (alternative PFAS). (**b**) The relative fluorescence signals were measured when different legacy PFAS (100 µM) were added to compete with the FABP bound fluorescence probes; 1,8-ANS were used for FABP 1, 2, 3, 4, 5, and FABP7, and bisANS was used for FABP6. Dose-response curves showing the competition between increasing compound concentrations and fluorescent probes for binding to the seven FABPs. The three compounds are displayed as follows: PFO3TDA(**c**) and HFPO-TeA(**d**). PFO3TDA (CAS, 330562-41-9) was tested with six concentrations (0.37, 1.11, 3.33, 10, 30, 50 µM) across seven FABPs; HFPO-TeA (CAS, 65294–16–8) was tested with six concentrations (3.12, 6.25, 12.5, 25, 50 100 µM) across seven FABPs. (**e**) PFO3TDA co-crystalized with FABP7 (PDB: 9NIU). The carboxylate head group of PFO3TDA formed two hydrogen bonds with residues arginine 127 and tyrosine 129. (**f**) The Gibbs free energy difference between ‘trans’ and ‘gauche’ conformations for PFOA, PFO3TDA (CF2-X-X-CF2), and PFO3TDA (O-CF2-CF2-X). (**g**) The diagram showing the interactions of endogenous ligands^29^ (oleic acid, retinoid acid and bile acid), legacy PFAS, and alternative PFAS (long-chain PFECAs) with FABPs and their corresponding peroxisome proliferator-activated receptors (PPARs), distributed across different body tissues^29,70–75^.

### Molecular basis for the selective binding of PFO3TDA to FABP7

To understand the mechanism underlying selective binding, we selected PFO3TDA and FABP7 for X-ray co-crystallization, as this interaction exhibited the strongest affinity among those tested in fluorescence displacement assays (K_D_ = 0.02 µM) (Table S2). The co-crystal structure (1.9 Å) revealed that PFO3TDA adopts a U-shape conformation within the binding pocket (Fig. 4d). However, the precise ligand conformation was only partially resolved due to the ambiguous electron density map around the ligand, likely a result of its inherent flexibility within the binding site. Despite this limitation, the U-shape conformation is clearly distinct from that of the endogenous ligand, palmitic acid, which was previously co-crystallized with FABP7 (PDB: 7E25)^52^. Analysis of the electron density map suggests that the observed U-shaped conformation is stabilized by a conformational shift in the ether linkages from the ‘trans’ to the ‘gauche’ conformation, allowing for a better fit within the binding pocket and enhanced interactions with surrounding hydrophobic residues. Additionally, the hydrogen bonds between the carboxylic acid group of PFO3TDA and residues R127 and Y129 further stabilizes the interaction, reinforcing its accommodation in FABP7.

To better understand the ligand’s conformation within the binding pocket, given the limited resolution of the crystallization results, we assessed the energetic cost of its conformational change. Density Functional Theory (DFT) calculations were performed to evaluate the torsional energy of PFO3TDA and PFOA (Fig. 4f). The energy changes for the CF2–X–X–CF2 dihedral angles were comparable between the two compounds, ranging from 0.52 to 6.68 kcal/mol for PFOA and 0.71 to 4.81 kcal/mol for PFO3TDA. However, the energy changes associated with the O–CF2–CF2– X dihedral angles were significantly lower for PFO3TDA (0.01-0.34 kcal/mol), suggesting that the ether linkage in PFECAs enhances conformational flexibility. Collectively, the co-crystal structure and DFT calculations demonstrate that the enhanced conformational flexibility of PFECAs allows them to better adapt to the binding pockets of FABPs, contributing to their selective binding.

## Discussion

We benchmarked the AS-MS platform to identify novel protein targets of prioritized chemical contaminants, with a 100% detection rate for strong-affinity ligands. Out of 24,050 assessed interactions (50 proteins × 481 compounds), AS-MS identified 35 compound-protein interactions. Among the 35 identified interactions, 27 were previously unreported, including 16 FABP-ligand interactions with high fold-change enrichment, which were successfully validated using a fluorescence displacement assay. Compared to the HTS bioassays used for the ToxCast program, our AS-MS platform has two major advantages: 1) it enables screening of a wide variety of proteins including novel proteins for which bioassays are not available, facilitating protein target scouting for environmental compounds at the proteome-wide level; 2) its throughput is much higher because thousands of compounds can be assayed simultaneously. Identifying protein targets often unlocks exciting research opportunities, but this process can be lengthy and challenging. For example, it took approximately 40 years to identify CRBN as the target of thalidomide^6^ and 30 years to discover aryl hydrocarbon receptor (AhR) as the target of dioxins^53^. Our AS-MS approach significantly accelerates this process by enabling scalable target discovery at both the protein and chemical levels. At the current rate, the screening of all ToxCast chemicals can be completed for 50 proteins within one week, and ∼1,500 proteins within one year, when purified His-tagged proteins are available. While production of proteins is a major limiting factor, our team at Structural Genomics Consortium (SGC) are the key contributors to the Target2035 program^54^ which seeks identify a chemical probe to every human protein by 2035. This ambitious program necessitates the purification of thousands of proteins, which could in turn also be used to perform ToxCast-like screens. We believe this will be a unique opportunity for target discovery for chemical contaminants approaching a proteome-wide level. However, like any technology, the AS-MS platform has its limitations: 1) it assesses protein bindings only, and additional functional experiments are needed to confirm functional consequences; 2) good MS sensitivity is essential for AS-MS screening, and thus complementary MS (*e.g.,* GC-MS) is needed to increase the coverage which might sacrifice throughput.

Of the 50 human proteins assessed beyond the scope of the ∼300 proteins covered by the ToxCast program, AS-MS binding was detected for 14 of them. This highlights the need for proteome-wide discovery of novel protein targets to achieve comprehensive insights into toxicity mechanisms. Particularly, the novel interactions discovered in this study for PFAS ‘forever chemicals’ revealed unexpected chemotype and structural selectivity. PFECAs were found to bind all 7 FABPs due to their more flexible conformation, whereas legacy PFAS bound only to FABP1. FABP1 has been well documented as a key regulator of the toxicokinetics of legacy PFAS^55^, which can disrupt its fatty acid transport function and hinder delivery to PPARs, a pathway well known to be linked to metabolic dysregulation and obesity^35,56,57^. Unfortunately, the broader FABP binding spectrum of PFECAs suggests an even greater toxicological impact by interrupting the FABP-PPAR pathways in multiple tissues beyond liver (Fig. 4g). Indeed, previous studies reported a stronger hepatic accumulation and toxicity of HFPO-TA compared to PFOA,^61,62^ consistent with its stronger binding affinity to liver FABP (FABP1). Collectively, this raises concerns that alternative PFAS may be regrettable substitutes for legacy compounds. Future studies are needed to evaluate their potential toxicity through the newly identified FABP pathways.

## Methods

### Chemical and reagents

The 1,040 ToxCast and 74 PFAS compounds were gifts from Dr. Keith A. Houck from the U.S. Environmental Protection Agency (EPA) (Table S5). The PFECAs include perfluoro-2,5,8-trimethyl-3,6,9-trioxadodecanoic acid (HFPO-TeA), 2,3,3,3-tetrafluoro-2-(heptafluoropropoxy)propanoic acid (HFPO-DA), perfluoro-2,5-dimethyl-3,6-dioxanonanoic acid (HFPO-TA), perfluoro-3,6,9-trioxatridecanoic acid (PFO3TDA), perfluoro(4-methoxybutanoic) acid (PFMOBA), and perfluoro-3,6-dioxaheptanoic acid (PFO2HpA), in which the first three were purchased from the Toronto Research Chemicals (Toronto, ON, CA), while the others were provided by the Evotec SE (Branford, CT) under contract to the U. S. Environmental Protection Agency (EPA) through a Material Transfer Agreement. The legacy PFAS include perfluorononanoic acid (PFNA), n-ethylperfluorooctanesulfonamide (N-EtFOSA), perfluorooctanesulfonic acid (PFOS), perfluorooctanesulfonamide (PFOSA), perfluorooctanoic acid (PFOA), and perfluorohexanesulfonate (PFHxS), all of which were provided by the Evotec SE (Branford, CT). Human FABP cDNAs were purchased from Origene, including FABP1 (NM_001443), FABP2 (NM_000134), FABP3 (NM_004102), FABP4 (NM_001442), FABP5 (NM_001444), FABP6 (NM_001445), and FABP7 (NM_001446). pET28-MHL vector was kindly provided by Structural Genomics Consortium (SGC). BL21(DE3) competent Escherichia coli was purchased from BioLabs Inc. (Whitby, ON, CA). 1-anilinonaphthalene-8-sulfonic acid (1,8-ANS), 4,4′-dianilino-1,1′-binaphthyl-5,5′-disulfonic acid dipotassium salt (bisANS) and Ni-NTA Agarose were purchased from Thermo Fisher Scientific (Ottawa, ON, CA). Desoxycholic acid (CAS 83-44-3) is one types of bile acids purchased from Toronto Research Chemicals Inc (Toronto, ON, CA). Methanol (HPLC grade), ultrapure water (HPLC grade), acetonitrile (HPLC grade), DMSO, glycerol, and formic acid were purchased from Fisher Scientific (Ottawa, ON, CA). HIS-Select nickel magnetic agarose beads and Triton X-100 were obtained from Sigma-Aldrich (St. Louis, MO, USA). Tris base, sodium chloride (NaCl), Tris(2-Carboxyethyl)phosphine (TCEP) were purchased from BioShop Canada Inc. (Burlington, ON, CA). Immidazole was provided by Bio Basic Canada Inc. (Markham, ON, CA).

### Protein Overexpression and Purification

Yang et al. have previously described the FABP protein overexpression process^25^. In brief, each individual human FABP was expressed in BL21 (DE3) *E. coli* cells as TEV-cleavable 6×histidine-tagged fusion proteins. BL21-Codon Plus cells were heat-shock transformed with isopropyl β-D-1-thiogalactopyranoside (IPTG)-inducible His-tagged FABPs ligand-binding domain (LBD) constructs. The transformed *E. coli* was plated on kanamycin-antibiotic plates and incubated at 37 °C overnight. The single colony of each plate was inoculated into LB media for incubation overnight, followed by overexpression by the addition of 1 mM IPTG. The bacteria cells were lysed by sonication in the lysis buffer as described before^25^. The FABP proteins were affinity-purified with Ni-NTA agarose according to the procedures from Cronet et al^63^.

Different from our previous study^25^, size exclusion fast protein liquid chromatography (FPLC) (HiLoad 16/60 Superdex 200, GE Healthcare, Life Sciences) was used to further purify the protein with 150 mM NaCl, 50 mM HEPES (pH 8.2), and 0.5 mM DTT, at a flow rate of 1 ml/min. The fractions of the elution were concentrated using Amicon centrifugal filters (10K, Sigma-Aldrich) and reconstituted with the stocking buffer (50 mM tri-HCl, 150 mM NaCl, 1 mM dithiothreitol, pH 8.0). The further purification was conducted because the impurities were found to impact the K_D_ values of 1,8-ANS binding to FABP1 by ∼5 times. Among 50 proteins, the information of 34 proteins was reported in our previous study^9^, the protein purification information of the rest of proteins was provided in the supporting information (Table S4).

### AS-MS platform

The prioritizing library is composed of 1,114 compounds, including 1,040 ToxCast chemicals and 74 per-and polyfluoroalkyl substances (PFAS). All 1,114 compounds were mixed together into one pool with final concentration of 10 μM of each compound. We selected 50 His-tagged human proteins (or domains thereof) from the SGC’s collection (Table S3), using approximately 0.03mg of each protein divided into two batches on two 96-well plates. The chemical pool was exposed to all 55 proteins with 3 replicates with the binding buffer (150 mM NaCl, 50 mM Tris-HCl, 0.1 mM TCEP, 0.5% glycerol, 0.01% Triton X-100, pH 7.5). 5 µL of Ni-NTA magnetic beads was added to the binding buffer, then adjusted to a final volume of 250 µL per sample. The final concentrations were set to 1 µM for the proteins and 100 nM for each chemical. The compounds and proteins were incubated on a rotary mixer at 4 °C for 30 minutes. After incubation, the 96-well plate was placed on a magnetic plate to immobilize the beads to help to remove the binding buffer. The beads were then washed twice with 100 µL of washing buffer 1 (150 mM NaCl, 50 mM Tris-HCl, 0.1 mM TCEP, 0.5% glycerol, 0.01% Triton X-100, 5 mM imidazole, and pH 7.5), followed by one wash with 100 µL of washing buffer 2 (150 mM NaCl, 50 mM Tris-HCl, 5 mM imidazole, and pH 7.5). When the beads were washed with washing buffer 2, they were resuspended and transferred to a new plate. After removing the washing buffer 2, 60 µL of methanol was added to each well to denature proteins and release chemical ligands. 50 µL of methanol extracts was aliquoted from each well and directly subjected to LC-MS analysis.

### Identification of protein hits using LC-MS

The AS-MS samples were analyzed using an Q Exactive Orbitrap mass spectrometer equipped with a Vanquish UHPLC system (Thermo Fisher Scientific, CA, USA). Chromatographic separation was conducted on an Accucore Vanquish C18 column (50 mm 2.1 mm × 1.5 μm, Thermo Scientific) with a flow rate of 0.2 mL/min. 1 µL of each sample was injected for analysis.

Two sets of mobile phases were used to facilitate the compound ionization: (1) Ultrapure water with 0.1% formic acid (A) and methanol with 0.1% formic acid (B) were used as the one set of mobile phases; and (2) Ultrapure water with 1 mM ammonium formate, pH 8 (A) and pure methanol (B) were used as the other set of mobile phases. The separation gradient elution started with 5% B for 1 minute, increased to 80% at 3 min, then reached to 100% at 4 min, held for 4 minutes, finally returned to 5% B over 1 min and equilibrated for 1 minute. The column temperature was maintained at 40 °C, and the sample compartment at 4 °C. Data acquisition was performed in full MS^1^ scan mode (100 - 800 *m/z*) with a resolution of R = 70,000 (at *m/z* 200) and a maximum of 3×10^6^ ions collected within 160 ms, in both positive and negative ionization modes.

### Fluorescence Displacement Assay

The binding of PFAS to recombinant FABPs was determined by the displacement of 1,8-ANS from proteins as described in our previous study except for FABP6^25^. Bis-ANS was used to determine the binding affinities of PFAS with FABP6 because of its stronger electrostatic and hydrophobic interactions with amino acid residues within the binding pocket of FABP6^64,65^.

The interactions between the legacy PFAS and FABPs (three replicates) were determined by displacing 1,8-ANS or bisANS by the PFAS from FABPs. For 1,8-ANS fluorescence probe, 40 μM of 1,8-ANS was incubated with 1 μM of each FABP for 30 min, followed by adding a fixed concentration (100 μM) of legacy PFAS individually to the mixture (n = 3). After another 30 min incubation, the fluorescence signal was measured by an Infinite 200 PRO multimode plate reader (Tecan Group Ltd., Switzerland) with 350 nm excitation wavelength and emission spectra between 420 nm and 600 nm. The same assay was used to determine the binding affinities for the PFECAs. Before determining the dissociation constant (K_D_) of PFECAs, the K_D_ of 1,8-ANS to different FABPs was required. These K_D_’s were experimentally derived from measuring maximal fluorescence intensities of bound 1,8-ANS at various concentrations (1.875-60 µM, n = 3) with different FABPs. The one-site binding model based in the Scatchard equation was fit to the displacement of protein-bound 1,8-ANS to calculate K_D_ to each FABP. Then, the K_D_ of six PFECAs was determined from the decreasing fluorescence signal induced by the displacement of 1,8-ANS from the FABP. Two concentration gradients were selected for the six PFECAs; the higher concentration gradient was selected for HFPO-TeA, HFPO-TA and HFPO-DA (0.41, 1.23, 3.70, 11.11, 33.33, and 100 µM); the lower concentration gradient was for PFO3TDA, PFO2HpA and PFMOBA (0.37, 1.11, 3.33, 10, 30, and 50 µM). Like the legacy PFAS, the different concentrations of PFECAs were added into 1,8-ANS (40 µM) and FABP (1 µM) mixture and the inhibition dose-response curves were plotted based on the relative fluorescence change. The K_D_ of PFAS to seven FABPs were determined through fitting to a sigmoidal model. The calculation method was as same as used in our previous study^25^.

The process for bisANS was similar to 1,8-ANS except for the determination of K_D_ value of the fluorescence probe. Six concentrations of FABP6 with 3-fold serial dilution (1.23, 3.7, 11.1, 33.3, 100, and 300 μM) were added into 1 μM of bisANS individually to determine the fluorescence signal intensity produced when the probes bind to more proteins. The fluorescence signal was also measured by an Infinite 200 PRO multimode plate reader with 390 nm excitation wavelength and emission spectra between 420 nm and 600 nm. Regression analysis was used to calculate the K_D_ of bisANS^64^. To determine the binding affinities of compounds to FABP6, the same concentration gradients of compounds were added to compete with 1 μM of bisANS bound to 10 μM of protein. The calculation steps were the same as 1,8-ANS. Principal component analysis (PCA) was conducted to analyze the FABP-ligand interaction behaviors based on the K_D_ values measured by the fluorescence displacement assay. The K_D_ of oleic acid (OA) to seven FABPs were obtained from literature^29,66,67^. Before analysis, the K_D_ values were transformed to-log_10_(K_D_) to ensure that higher values indicate stronger protein-ligand interactions. In PCA biplot, the blue dots represent different ligands and the arrows of FABPs reflect their contribution to the principal components.

### Calculating Protein Binding Cavity

The volume of the binding cavity of seven FABPs was obtained by calculating solvent-accessible (SA) volume of proteins through the Computed Atlas of Surface Topography of proteins (CASTp), an online resource that computes the geometric and topological properties of proteins^41^. The crystal structures of proteins were downloaded from Protein Data Bank, with the resolution of X-ray diffraction less than 2.0 Å were uploaded to CASTp to measure the SA volume. The radius of the probe was set to 1.4 Å as a default. The selected protein structure IDs with the measured SA volumes are listed in SI (Table S1).

### Protein Crystallization

Perfluoro-3,6,9-trioxatridecanoic acid (PFO3TDA) was co-crystallized with FABP7 protein. A mixture of 2 mg/mL of compound and 9.7 mg/mL of TEV-cleavable 6×histidine-tagged FABP7 was prepared in a crystallization buffer containing 12.5% (w/v) PEG 1000, 12.5% (w/v) PEG 3350, 12.5% (v/v) MPD, 0.03 M of each halide, 0.1 M MES/imidazole and pH 6.5. The protein-ligand mixture was mixed with the crystallization buffer in a 1:1 ratio, with 200 nL of each component, and set up in a sitting-drop crystallization plate at 20 °C. The crystal was harvested and flash-cooled in liquid nitrogen without additional cryoprotectant. Diffraction data were collected using a Rigaku FR-E SuperBright source. The data were processed and scaled using HKL-3000, and the structure was solved by molecular replacement in PHASER using the starting coordinates 6L9O. Refinement was performed using BUSTER (v2.10.3). The final coordinates of the protein and ligand were deposited in the PDB under accession code 9NIU.

### DFT Calculations

The changes of torsional energies with bond angles of legacy PFAS (PFOA) and PFECAs (PFO3TdA) were calculated using the GAUSSIAN 16 software. The molecular geometries of PFO3TDA and PFOA were optimized using the B3LYP functionals and 6-311G basis set. No solvent correction was applied to best emulate the solvent-free conditions of the hydrophobic binding pockets of FABP proteins. The geometry-optimized structures were used to perform bond angle scans of CF2-O-CF2 in PFO3TDA and CF2-CF2-CF2 in PFOA with a step size of 15o. Similarly, dihedral angle scans were performed for CF2-CF2-CF2-CF2 in PFO3TDA and PFOA, as well as O-CF2-CF2-X in PFO3TDA, with a step change of 10° (Fig. S6, Fig. S7).

## Code availability

The source code and demo datasets can be found via GitHub at https://github.com/huiUofT/EAS-MS.

## Acknowledgements

We extend our deepest gratitude to the Structural Genomics Consortium-Toronto (SGC-Toronto) and Structural Genomics Consortium-Frankfurt (SGC-Frankfurt) for their generous contribution of proteins, which were invaluable to this study. From SGC-Toronto, we acknowledge the provision of GID4, SLC9C1 and C2, DNMT3A, SKP1, SKP1FBXO22, MSH6, USP37, SETDB1, HTT/HAP40, WDR55, SEC31A, DCAF7, HDAC6, CRBN, TLE4, DDB1, COPB2, WDR5, FBXW7, RFWD3, DCAF1, WDR91, LRRK2, HAT1, PIAS1, DDX1, HUWE1WWE, YTHDF2, YTHDC1, CHIP, and USP21. From SGC-Frankfurt, we acknowledge the provision of 1TIF1A, 2TIF1A, TRIM28, TRIM7, TRIM21, BIR1, BUCK, and SPSB2. We also appreciate Wyatt Yue from Newcastle University for providing AASSLKR and AASSSDH proteins. We are especially grateful to Elisa Gibson, Pegah Ghiabi, Carla Kharadjian, Serah Kimani, Renu Chandrasekaran, Taraneh Hajian, Andreas C. Joerger, Krishna Saxena, Yeojin Kim, and Madhushika Silva for their dedicated efforts in protein preparation. Additionally, we sincerely acknowledge Albina Bolotokova, Vijayaratnam Santhakumar, and Suzanne Ackloo for their invaluable contributions in preparing benchmarking chemicals and coordinating the delivery of chemicals and proteins. We also extend our gratitude to Keith A. Houck from the U.S. Environmental Protection Agency (EPA) for providing ToxCast and PFAS compound data.

Furthermore, we would like to thank Dafydd Owen (Pfizer), Judith Guenther (Bayer), Scott Johnson (Bristol Myers Squibb), Oliver Kraemer (Boehringer Ingelheim), Julian Schmid (Merck), Juncai Meng (Janssen), Lawrence Szewczuk (Janssen), Xidong Feng (Pfizer), Wenyi Hua (Pfizer), Anja Giese (Bayer), Matteo Aldeghi (Bayer), Nidhi Arora (Takeda), James Kiefer (Genentech), Ingo Hartung (Merck), and Matthew Troutman (Pfizer) for their valuable advice and discussions on method development. The Structural Genomics Consortium is a registered charity (no. 1097737) that receives funding from Bayer AG, Boehringer Ingelheim, Bristol Myers Squibb, Genentech, the EU/EFPIA/OICR/McGill/KTH/Diamond Innovative Medicines Initiative 2 Joint Undertaking ([EUbOPEN grant 875510]), Janssen, Pfizer, and Takeda. This research was supported by a Natural Sciences and Engineering Research Council of Canada (NSERC) Discovery Grant, an Ontario Early Researcher Award, the Canadian Institutes of Health Research (CIHR), and the National Institutes of Health (NIH) (U54AG065187). The authors also acknowledge the support of instrumentation grants from the Canada Foundation for Innovation, the Ontario Research Fund, and an NSERC Research Tools and Instruments Grant. AME holds the Temerty Nexus Chair of Health Technology and Innovation at the University of Toronto.

## Author contributions

DY, XW, JL, JS, YG: AS-MS method development DY, XW, XQ, CC: Chemical library screening

RJH, NB, TSB, HZ, DLB, NGB: Proteins preparation and coordination

PN, DS: DFT calculation

HZ, AD: Protein Crystallization

HP, LH, HK, MLD, AME, CHA: Supervision, review and editing

DY, XW, HP: Writing, reviewing and editing with input from all

