## Supporting Information for "High-throughput protein target mapping enables accelerated bioactivity discovery for ToxCast and PFAS compounds"

**Table of contents**

P.3 – Supplementary Fig. S1 | Benchmark the AS-MS platform using 8 human proteins and their positive ligands.

P.4 – Supplementary Fig. S2 | SDS-PAGE and western blot results of seven purified His-tagged human FABPs.

P.5 – Supplementary Fig. S3 | Scaffolds of compounds involved in the 35 detected protein-ligand interactions.

P.6 – Supplementary Fig. S4 | The fluorescence intensities measured to calculate the $K_D$ of the fluorescence probes to each FABP.

P.7 – Supplementary Fig. S5 | Lack of Interaction Between Short-Chain PFECAs and FABPs.

P.8 – Supplementary Fig. S6 | The conformations of palmitic acid in FABPs with the Newman projections to indicate the bond rotations.

P.9 – Supplementary Table S1 | The PDB I.D. of the crystal structures of FABPs used for calculating the solvent-accessible volume through the CASTp.

P.10 – Supplementary Table S2 | The information of the hits to target proteins detected by the ASMS platform.

P.11-13 – Supplementary Table S3 | Information of 50 human proteins screening in the AS-MS platform.

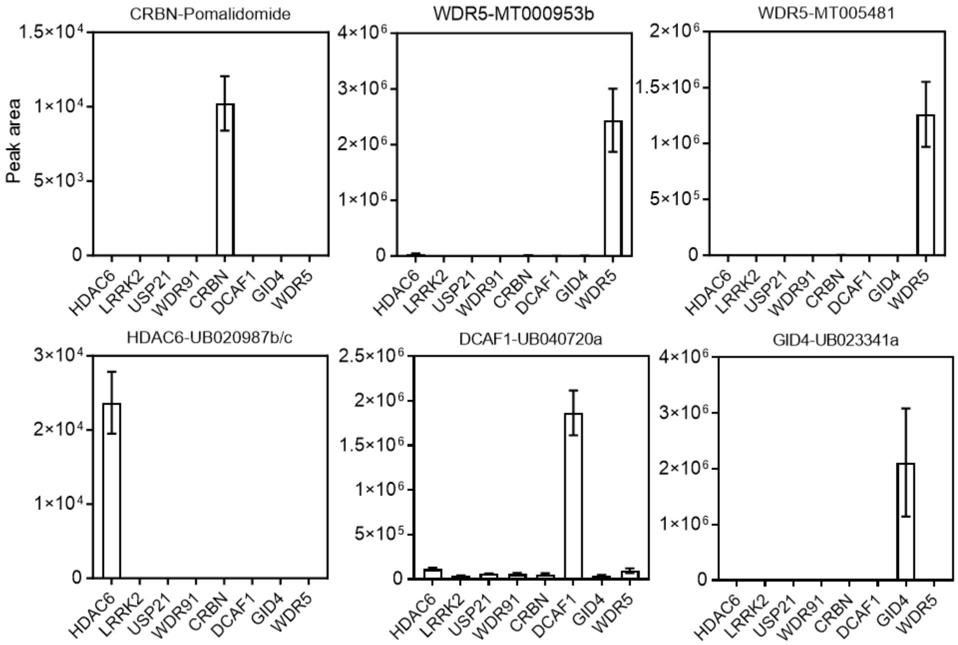


**Figure S1**. **Benchmark the AS-MS platform using 8 human proteins and their positive ligands.** The platform successfully identified 6 known ligands binding to their corresponding proteins from a chemical library of 101 compounds. The proteins tested include histone deacetylase 6 (HDAC6), leucine-rich repeat serine/threonine-protein kinase 2-WDR domain (LRRK2), ubiquitin carboxyl-terminal hydrolase 21 (USP21), WD repeat-containing protein 91 (WDR91), cereblon (CRBN), DDB1- and CUL4-associated factor 1-WDR domain (DCAF1), glucose-induced degradation protein 4 homolog (GID4), and WD repeat-containing protein 5 (WDR5).

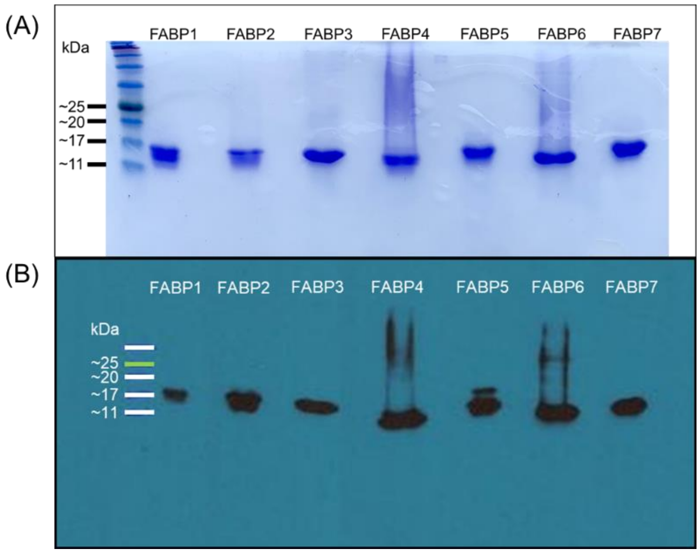


**Figure S2**. **SDS-PAGE and western blot results of seven purified His-tagged human FABPs**. (A) Coomassie blue stained SDS-PAGE indicated the purified human FABPs. (B) The overexpressed His-tagged FABPs in *Escherichia coli* were detected by the His-tagged antibody through western blot.

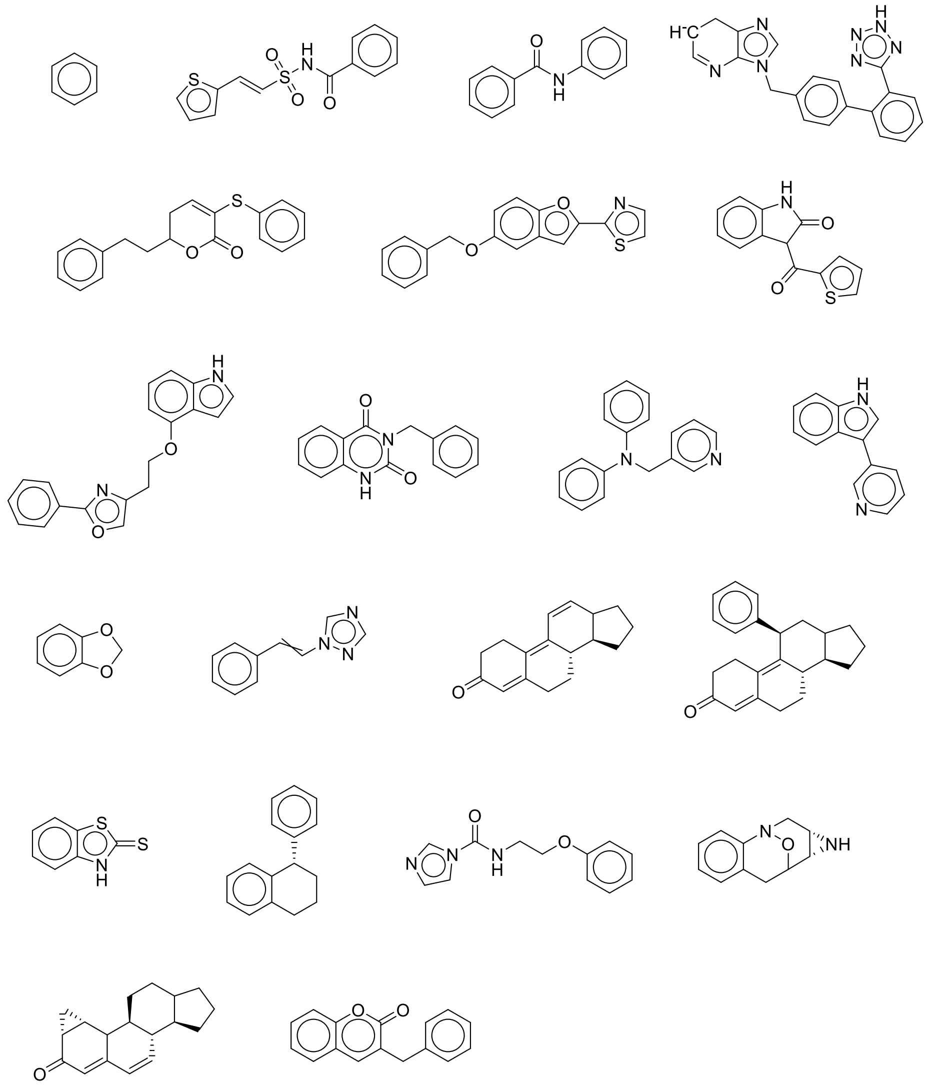

**Figure S3. Scaffolds of compounds involved in the 35 detected protein-ligand interactions.** A total of 21 distinct scaffolds were identified, while 12 compounds had unknown scaffolds.

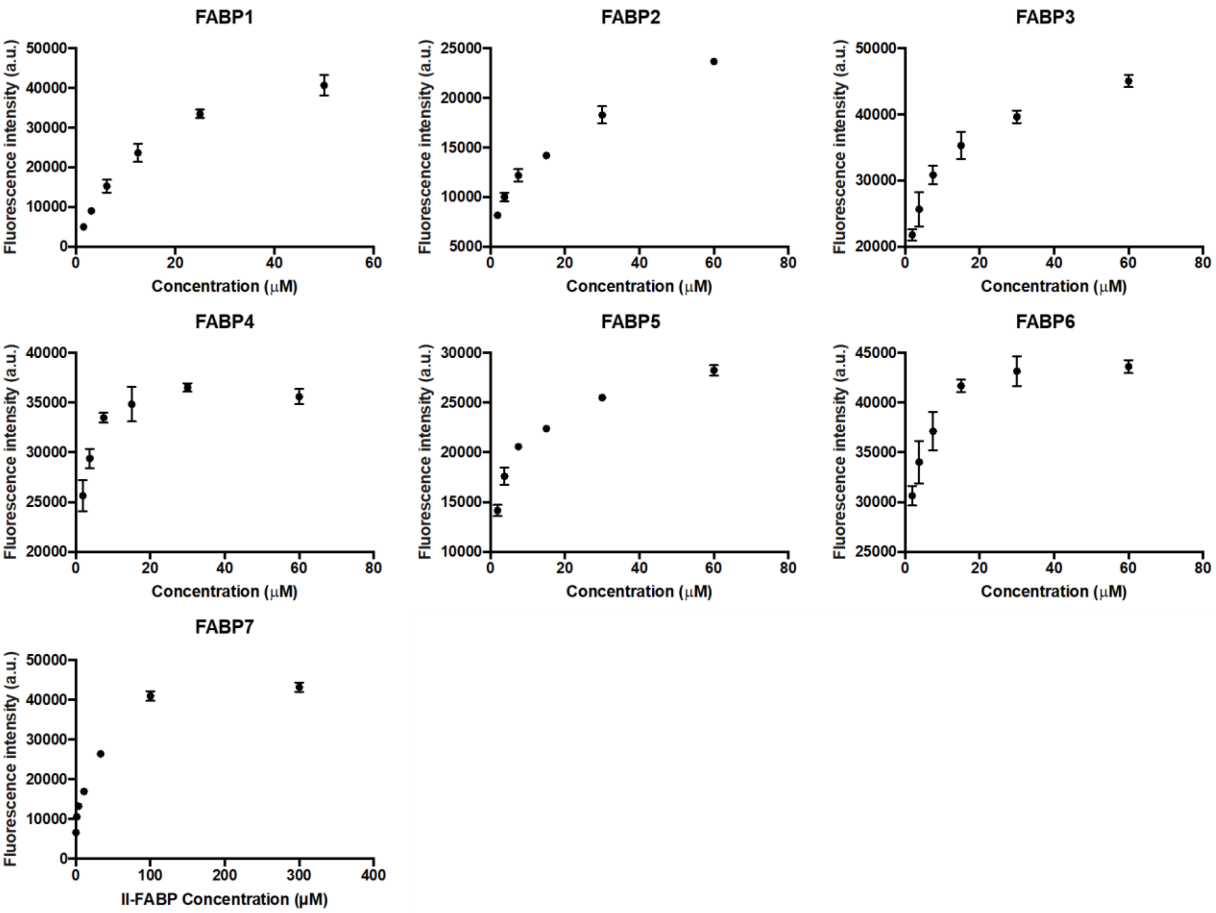


**Figure S4**. **The fluorescence intensities measured to calculate the $K_D$ of the fluorescence probes to each FABP.** The fluorescence signal was measured when the increasing concentration of 1,8-ANS added into 1 μM of FABP1, 2, 3, 4, 5, 6. While the signal of bisANS was measured when the increasing amount of Il-FABP added into 1 μM of bisANS.

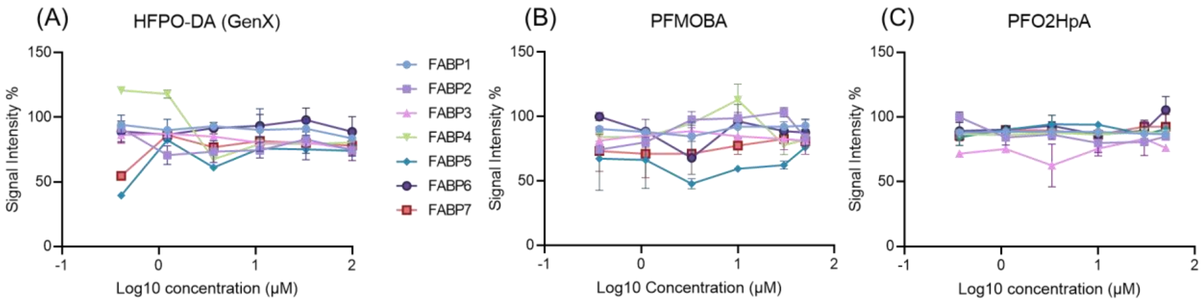


**Figure S5. Lack of Interaction Between Short-Chain PFECAs and FABPs.** Fluorescence measurements were conducted by adding varying concentrations of PFECAs to a mixture containing 1 µM protein and 40 µM 1,8-ANS. Dose-response curves indicate no competitive binding between the fluorescence probe and HFPO-DA (A), PFMOBA (B), or PFO2HpA (C) across seven FABPs. HFPO-DA was tested at six concentrations: 0.41, 1.23, 3.70, 11.1, 33.3, and 100 µM, while slightly lower concentrations were used for PFMOBA and PFO2HpA: 0.37, 1.11, 3.33, 10, 30, and 50 µM. 1,8-ANS functioned as the fluorescence probe for FABPs 1, 2, 3, 4, 5, and 7, whereas bis-ANS was used for FABP6.

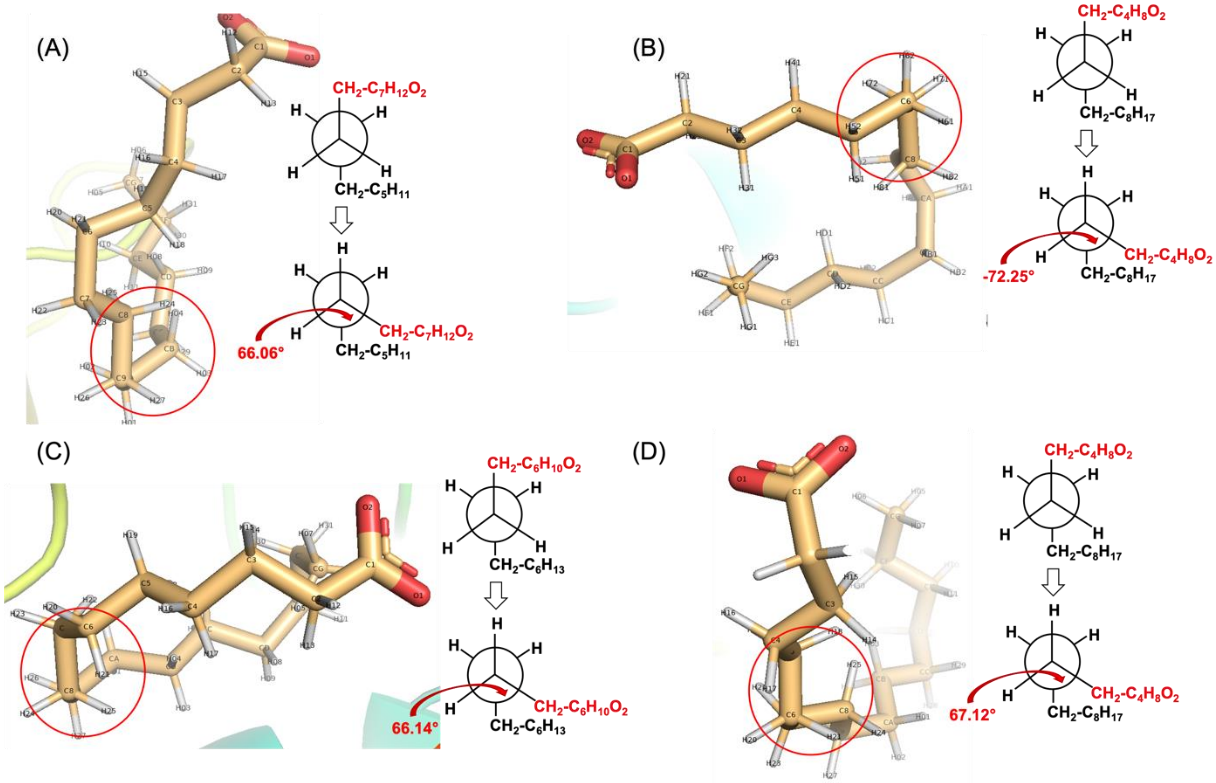


**Figure S6. The conformations of palmitic acid in FABPs with the Newman projections to indicate the bond rotations.** The crystal structures of FABPs with palmitic acid as the original ligands were used to determine the bond rotations of ligands in the proteins. The protein structures include (A) FABP1 (3STM), (B) FABP3 (6AQ1), (C) FABP4 (2HNX) and (D) FABP5 (1B56).

**Table S1**. **The PDB I.D. of the crystal structures of FABPs used for calculating the solvent-accessible volume through the CASTp.**

| Protein | PDB ID | Solvent-Accessible volume ($Å^3$) |
|---|---|---|
| FABP1 | 3STM | 361 |
| FABP2 | 3AKM | 288 |
| FABP3 | 3WBG | 226 |
| | 6AQ1 | 260 |
| | 4WBK | 258 |
| | 3RSW | 399 |
| | 5B28 | 285 |
| | 5HZ9 | 237 |
| | 4TJZ | 246 |
| | 3WXQ | 269 |
| | 4TKH | 250 |
| | 3WVM | 256 |
| FABP4 | 5y12 | 263 |
| | 3Q6L | 257 |
| | 6AYL | 411 |
| | 2NNQ | 325 |
| FABP5 | 5HZ5 | 256 |
| | 4LKP | 218 |
| | 5UR9 | 353 |
| | 1B56 | 311 |
| FABP6 | 1O1V | 377 |
| | 5L8I | 383 |
| | 2MM3 | 420 |
| FABP7 | 7E25 | 272 |
| | 6L9O | 298 |
| | 1FE3 | 323 |
| | 1FDQ | 295 |
| | 5URA | 333 |

**Table S2**. **The information of the hits to target proteins detected by the ASMS platform.** The table includes CAS number, fold changed compared to the other proteins, corresponding *p values* and measured dissociation constants ($K_D$ μM) using fluorescence displacement assays of the discovered ligands.

| **Proteins** | **CAS** | **ASMS Fold change** | ***p value*** | **KD (μM)** |
|---|---|---|---|---|
| FABP1 | 135080-03-4 | 10.06 | 1.14E-02 | 4.45 |
| FABP1 | 136042-19-8 | 20.34 | 1.51E-02 | 0.56 |
| FABP1 | 149413-74-1 | 10.59 | 1.41E-03 | |
| FABP1 | 1763-23-1 | 53.10 | 3.53E-30 | 8.1 |
| FABP1 | 207736-05-8 | 20.29 | 1.32E-02 | 1.68 |
| FABP1 | 31506-32-8 | 74.91 | 3.07E-35 | |
| FABP1 | 335-67-1 | 87.09 | 2.45E-19 | 18.74 |
| FABP1 | 375-95-1 | 17.25 | 3.36E-04 | 5.4 |
| FABP1 | 3871-99-6 | 153.32 | 4.25E-34 | 11.19 |
| FABP1 | 4151-50-2 | 34.12 | 7.56E-33 | 16.58 |
| FABP1 | 50-65-7 | 22.00 | 6.29E-03 | 9.69 |
| FABP1 | 534-52-1 | 13.75 | 6.18E-03 | |
| FABP1 | 754-91-6 | 670.37 | 5.64E-37 | 5.87 |
| FABP1 | 88-85-7 | 59.14 | 1.07E-02 | |
| FABP1 | NOCAS_48518 | 48.29 | 2.00E-04 | 8.05 |
| FABP2 | 112733-06-9 | 20.05 | 9.11E-03 | 2.68 |
| FABP2 | 501027-49-2 | 32.51 | 2.57E-03 | 31.92 |
| FABP3 | 460081-99-6 | 22.89 | 5.02E-03 | |
| FABP4 | 120-83-2 | 56.95 | 3.84E-02 | |
| FABP4 | 162706-14-1 | 42.97 | 7.00E-03 | 3.26 |
| FABP4 | 22781-23-3 | 13.07 | 2.28E-02 | |
| FABP4 | 55406-53-6 | 18.37 | 2.99E-02 | |
| FABP6 | 83657-24-3 | 27.01 | 6.63E-03 | |
| FABP7 | 330562-41-9 | 53.95 | 6.57E-39 | 0.02 |
| SKP1 | 76578-14-8 | 14.40 | 7.47E-04 | |
| SKP1 | 77-89-4 | 10.81 | 7.23E-04 | |
| SKP1FBXO22 | 67747-09-5 | 22.04 | 1.89E-03 | |
| SKP1FBXO22 | NOCAS_48166 | 16.40 | 1.90E-03 | |
| WDR55 | 10161-33-8 | 60.32 | 5.10E-03 | |
| AASSLKR-GFP | 427-51-0 | 10.70 | | |
| BIR1 | 149-30-4 | 10.63 | 2.95E-02 | |
| CRBN | 84371-65-3 | 12.16 | 2.03E-02 | |
| DCAF7 | 291305-06-1 | 30.20 | 2.83E-02 | |

**Table S3. The information of 50 human proteins screening in the AS-MS platform.**

| Protein abbreviation | Full protein name | UniProt ID | Biological functions |
|---|---|---|---|
| DCAF1 | DDB1- and CUL4-associated factor 1-WDR domain | Q9Y4B6 | E3 ligase receptor protein |
| DNMT3A | DNA (cytosine-5)-methyltransferase 3A-PWWP domain | Q9Y6K1 | DNA methylation (histone methyllysine reader domain thereof) |
| LRRK2 | Leucine-rich repeat serine/threonine-protein kinase 2-WDR domain | Q5S007 | WD domain of LRRK2 proteins |
| RFWD3 | E3 ubiquitin-protein ligase RFWD3 | Q6PCD5 | E3 ligase receptor protein |
| COPB2 | Coatomer subunit beta-WDR domain 1 | P35606 | ER-Golgi transport |
| DDB1 | DNA damage-binding protein 1 | Q16531 | E3 ligase adaptor protein |
| WDR91 | WD repeat-containing protein 91 | A4D1P6 | Endosomal maturation |
| DDX1 | ATP-dependent RNA helicase DDX1 | Q92499 | RNA Helicase |
| SETDB1 | Histone-lysine N-methyltransferase SETDB1 | Q15047 | Histone binding domain of H3K9me3 methyltransferase |
| SEC31A | Protein transport protein Sec31A | O94979 | Protein transport |
| SKP1 | S-phase kinase-associated protein 1 | P63208 | E3 ligase adaptor protein |
| MSH6 | DNA mismatch repair protein Msh6 | P52701 | Component of the post-replicative DNA mismatch repair system (MMR) |
| FBXW7 | F-box/WD repeat-containing protein 7 | Q969H0 | E3 ligase receptor protein |
| CHIP | E3 ubiquitin-protein ligase CHIP | Q9UNE7 | E3 ubiquitin-protein ligase |
| PIAS1 | E3 SUMO-protein ligase PIAS1 | O75925 | E3-type small ubiquitin-like modifier (SUMO) ligase |
| 1TIF1A | Bromodomain of transcription intermediary factor 1-alpha | O15164 | Transcriptional coactivator |
| 2TIF1A | PHD and bromodomain of transcription intermediary factor 1-alpha | O15164 | Transcriptional coactivator |
| TRIM28 | Transcription intermediary factor 1-beta | Q13263 | Nuclear corepressor for KRAB domain-containing zinc finger proteins (KRAB-ZFPs) |
| TRIM21 | Tripartite motif-containing protein 21 | Q62191 | E3 ubiquitin-protein ligase |

| | | | |
|---|---|---|---|
| BIR1 | Baculoviral IAP repeat-containing protein 3 BIR1 domains | Q13489 | E3 ubiquitin-protein ligase |
| BUCR | Baculoviral IAP repeat-containing protein 3 BIR3-UBA-CARD-RING domains | Q13489 | E3 ubiquitin-protein ligase |
| TRIM7 | E3 ubiquitin-protein ligase TRIM7 | Q9C029 | E3 ubiquitin-protein ligase |
| SPSB2 | SPRY domain-containing SOCS box protein 2 | Q99619 | E3 ubiquitin-protein ligase |
| HAT1 | Histone acetyltransferase type B catalytic subunit | O14929 | Acetyltransferase |
| HTT/HAP40 | Huntingtin in complex with HAP40 | P42858/P 23610 | Mutated in Huntington's Disease |
| TLE4 | Transducing-like enhancer protein 4 | Q04727 | Transcriptional corepressor that binds to a number of transcription factors |
| WDR55 | WD repeat-containing protein 55 | Q9H6Y2 | Ribosomal maturation |
| SKP1FBXO22 | F-box only protein 22 | Q8NEZ5 | E3 ligase receptor protein |
| DCAF7 | DDB1- and CUL4-associated factor 7 | P61962 | E3 ligase receptor protein |
| YTHDC1 | YTH domain-containing protein 1 | Q96MU7 | Regulator of alternative splicing that specifically recognizes and binds N6-methyladenosine (m6A)-containing RNAs |
| YTHDF2 | YTH domain-containing family protein 2 | Q9Y5A9 | Specifically recognizes and binds N6-methyladenosine (m6A)-containing RNAs, and regulates their stability |
| AASSLKR | Alpha-aminoadipic semialdehyde synthase, lysine ketoglutarate reductase domain | Q9UDR5 | Bifunctional enzyme that catalyzes the first two steps in lysine degradation |
| AASSSDH | Alpha-aminoadipic semialdehyde synthase, saccharopine dehydrogenase | Q9UDR6 | Bifunctional enzyme that catalyzes the first two steps in lysine degradation |
| THEMIS | Thymocyte-expressed molecule involved in Selection CABIT domain | Q8N1K5 | Essential for T cell development and regulates phosphatases downstream of T cell receptor-associated kinases. |
| HUWE1WWE | WWE domain of HECT-type HUWE1 E3 ubiquitin ligase | Q7Z6Z7 | E3 ligase ADPR binding domain |
| SLC9C1 | Solute carrier family 9 member C1 | Q4G0N8 | Sperm-specific sodium/hydrogen exchanger involved in intracellular pH regulation of spermatozoa. |

| | | | |
|---|---|---|---|
| SLC9C2 | Solute carrier family 9 member C2 | Q5TAH2 | Involved in pH regulation |
| USP21 | Ubiquitin carboxyl-terminal hydrolase 21 | Q9UK80 | Deubiquitinases |
| WDR5 | WD repeat-containing protein 5 | P61964 | Involve in histone methyltransferase complexes |
| HDAC6 | Histone deacetylase 6 | Q9UBN7 | Deacetylase for non-histone proteins |
| GID4 | Glucose-induced degradation protein 4 homolog | Q8IVV7 | E3 ubiquitin-protein ligase |
| CRBN | Protein cereblon | Q96SW2 | E3 ubiquitin-protein ligase |
| FABP1 | Fatty acid-binding protein, liver | P07148 | Fatty acid (FA) transport, metabolism, and energy homeostasis |
| FABP2 | Fatty acid-binding protein, intestinal | P12104 | Dietary long-chain fatty acids absorption and transport, metabolism, and energy homeostasis |
| FABP3 | Fatty acid-binding protein, heart | P05413 | Fatty acid transport and β-oxidation in muscle and cardiac cells |
| FABP4 | Fatty acid-binding protein, adipocyte | P15090 | Fatty acid metabolism and inflammation regulation |
| FABP5 | Fatty acid-binding protein 5 | Q01469 | Fatty acid metabolism, inflammation regulation and epidermal barrier functions |
| FABP6 | Gastrotropin | P51161 | Bile acid transport and recycling |
| FABP7 | Fatty acid-binding protein, brain | O15540 | Fatty acid transport and metabolism, brain development |
| USP37 | Ubiquitin carboxyl-terminal hydrolase 37 | Q86T82 | Deubiquitinating enzyme |

97
